# Investigating Confidence Judgments using Reviews in Human and Monkeys

**DOI:** 10.1101/741561

**Authors:** Frederic M. Stoll, Emmanuel Procyk

## Abstract

Confidence judgments are self-assessments of the quality of one’s own performance, and are a crucial aspect of metacognitive abilities. The underlying neurobiological mechanisms are poorly understood. One approach to understanding these mechanisms would be to take advantage of putative metacognitive abilities in non-human models. However, many discrepancies exist between human and non-human studies on metacognition due to the mode of reporting judgements. We here present an attempt to directly compare human and non-human primates’ metacognitive abilities using a protocol assessing confidence judgments. After performing a categorization test, subjects could either validate their choice or review the test. We could assess whether subjects detected their errors and how they corrected them according to their confidence, and importantly did so in both human and non-human primates. 14 humans and 2 macaque monkeys were tested. Humans showed a well-adapted use of the review option by reviewing more after incorrect choices or difficult stimuli. Non-human primates did not demonstrate a convincing use of the review or validate opportunity. In both species, reviewing did not improve performance. This study shows that decisions to review under uncertainty are not naturally beneficial to performance and is rather perturbed by biases and alternative low-cognitive cost strategies.

## Introduction

After a decision, and before receiving any feedback, we may feel more or less confident that it was the correct one. For example, being confident you locked your car you will continue shopping, but if you are unsure, you will probably go back and check. Subjective confidence is one core aspect of metacognitive abilities, which represent higher order mental processes by which we monitor and control our own cognition^1^. In Humans, subjective confidence has generally been studied using prospective or retrospective questionnaires, requiring explicit verbal reports (e.g. confidence ratings^2,3^). Subjects are fairly good at judging their accuracy in both perceptual or mnemonic tasks^4^. Both prospective and retrospective confidence ratings appear to be highly correlated with one’s own performance, even if humans appear generally overconfident that their choice was/will be correct. Many theories have been proposed to explain how confidence judgments predict accuracy, but they do not account for the wide range of behavioural observations^5^. Nevertheless, an influential proposition is that people rely on inference to judge their performance, by accessing important features such as familiarity or difficulty with the test^6,7^.

To study confidence in non-human animals, researchers have adopted a broader view than for human studies, and explored a wider range of behaviour that might elicit metacognitive processes. Initial demonstration of animal metacognitive abilities involved “uncertain response” protocols, in which difficult tests could be avoided by the use of an alternative non-directed option. This option can be seen as a “choice to not choose”, and in theory it should be elicited by higher uncertainty about the outcome of the main choice^8,9^. Species including monkeys or dolphins efficiently used this option, although doubts have been raised concerning the involvement of metacognition in making such decisions^10^. Other studies attempted to approximate human protocols by assessing confidence judgment using a betting procedure^11^. In these tasks, monkeys were primarily asked to perform a perceptual test. After each trial they were then required to rate their confidence by ‘betting on their success’ (validating their choice) or alternatively by using a safe option. Monkeys correctly took the opportunity to bet, by betting more when correct^11,12^. Finally, a large body of work has focused on information-seeking behaviour as a mean of understanding metacognitive abilities in non-human animals^13,14^. This approach appeared more justified from an ecological point of view, as metacognition might enable animals to search for or verify information to improve their decisions. Such behaviour is thought to reflect higher cognitive processing. This is supported by studies highlighting that an animal will search for information when ignorant or uncertain about what to do^12–16^. Taken together, these studies show that information-seeking is targeted, ordered and optional, suggesting that such a search derives from metacognitive processes.

However, although information-seeking protocols have provided meaningful behavioural insight concerning putative non-human metacognitive abilities, they often appear inappropriate from a psychophysical or neurophysiological perspective, and are frequently criticised for poor control over experimental conditions. Notably, it has been suggested that simpler heuristics could be used to solve these tasks, questioning the need to rely on metacognition in such information-seeking paradigms^17^. Also, available paradigms did not assess an important feature of metacognition, which goes beyond simply validating or not a response based on uncertainty, to explicitly test the use of this decision to further seek information for improving performance. The objective of our current study was to test a new protocol adapted to human and non-human primates and devoted to study information-seeking and metacognition, possibly in the context of neurophysiological studies.

## Results

We designed a new behavioural task that allowed subjects to freely choose whether or not to review a test before validating their decisions (Figure 1A). In each trial, subjects were first (1) asked to report the angle (right or left) of an oriented grating and then (2) proposed to review or validate their choice. If subjects decided to review, they were able to go through the categorization and decision stages again. The aim of this task was to promote review depending on one’s own perceived uncertainty, as well as allowing subjects to use their uncertainty to guide further decisions. We first tested 14 human subjects in this new task. Subjects’ performances in the categorization were standardized using a staircase procedure prior to the experiment, defining the 3 levels of difficulty used thereafter (see Methods and Figure 1B).

**Figure 1.**
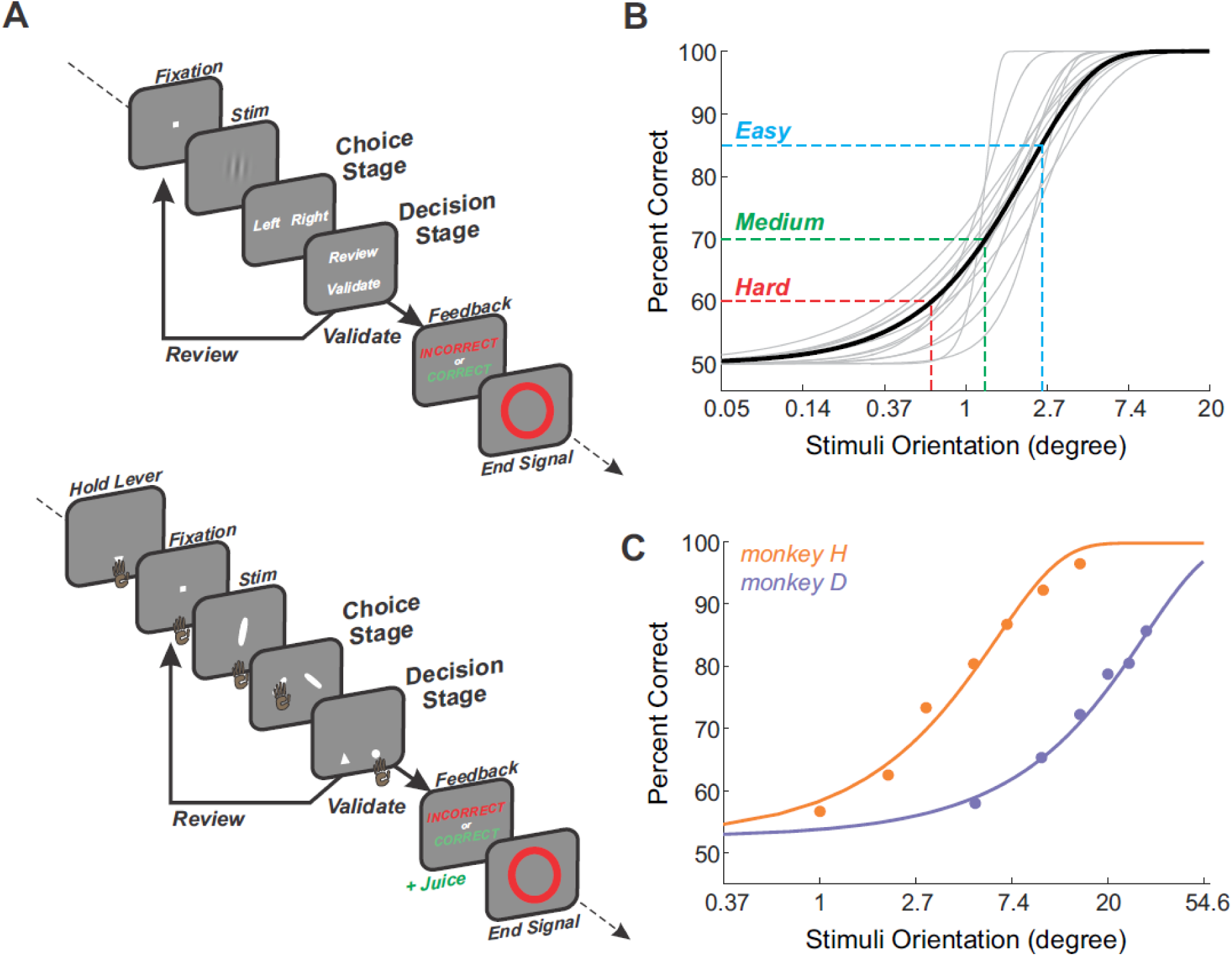
Confidence Judgment Task and categorization performance. **A**, The events of an individual trial of the task used for human subjects (top) and monkeys (bottom). Frames represent main successive events during a trial. Subjects had to choose whether the stimulus was oriented to the right or to the left. After the right/left choice (*Choice Stage*), subjects could decide whether to validate or review the stimulus to retry the test (*Decision Stage*). Visual feedback was given only after subjects’ decision to validate. In the monkey version of the task (bottom panel), they were required to hold a lever to initiate a trial and stimuli were oriented bars. Also, choice targets were represented by rightward and leftward oriented bars, randomly positioned. Decision targets were represented by a triangle (review option) or a circle (validate option), also randomly positioned. Additionally, correct trials were rewarded by juice, incorrect were penalized by a timeout. **B and C**, Individuals and average psychometric curve (binomial GLM using a logistic regression) showing performance across stimuli orientations (absolute value by pooling rightward and leftward stimuli) tested during the staircase procedure in human subjects (B) or across different sessions in both monkeys (C). Three absolute stimuli orientations were chosen from individual psychometric curves to elicit 60, 70 and 85% of correct choices (Hard, Medium and Easy conditions respectively).

### An adapted use of the review option in Humans

As expected, human subjects used the review option depending on categorization difficulty (Figure 2A). They reviewed significantly more often for harder stimuli than easier ones (mixed-effect *glm*, factor difficulty, F=33.6, p=3.3e-9). Contrary to our predictions, however, we observed no gain in performance after reviews (Figure 2B). In fact, the number of successive reviews a subject performed had a significantly detrimental effect on performance, notably after 2 successive reviews (mixed-effect *glm*; factor difficulty, F=61.7, p=4.0e-19; factor nbReview, F=4.24, p=0.016; the interaction did not survive model selection) (post hoc comparison of performance after 0 vs. 2 reviews, Wald test p=0.013; other conditions, p>0.12). Thus subjects took the opportunity to review when it made sense to do so, but gained no benefits on performance. This suggests that the probability to review depended on confidence level but that reviewing could not be leveraged to increase performance.

**Figure 2.**
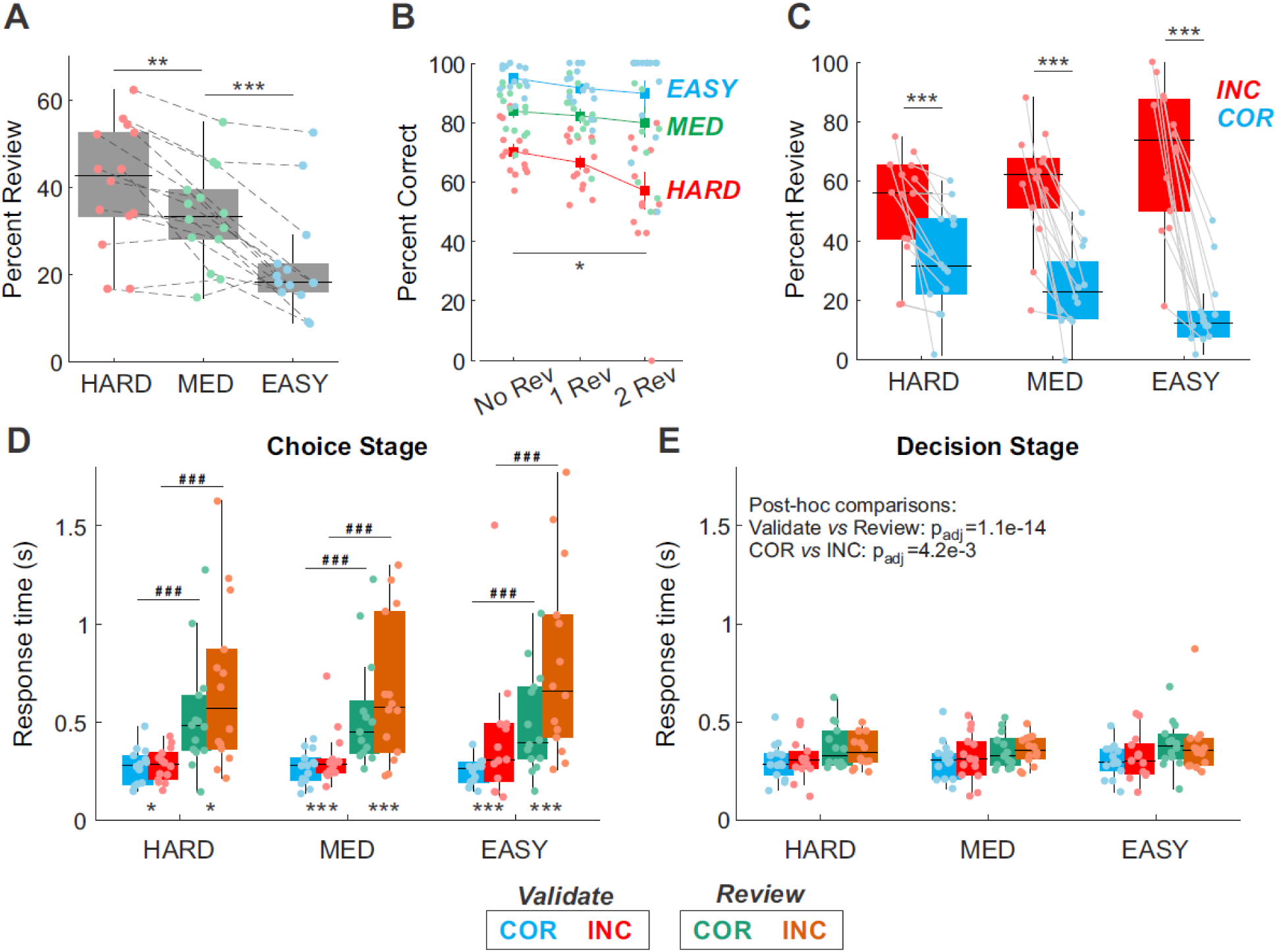
Behavioural performance of humans. **A**, Average percent of review choices for the three difficulty levels. **B**, Average final performance for no review (validate at the first decision), and after 1 or 2 successive reviews. **C**, Percent of reviews for each difficulty level separated by performance at the 1^st^ choice stage (either correct or incorrect). **D & E**, Median response times at the choice (**A**) and decision (**B**) stages depending on performance (correct or incorrect choice, blue/green and red/orange respectively) and difficulties. Blue/red colours represent validated trials, while green/orange colours reviewed ones. Significant post-hoc comparisons (Wald test with FDR correction) are reported as follow: *p<0.05; **p<0.01; ***p<0.001. In **D**, stars and sharp symbols stand respectively for a significant difference between incorrect/correct or validate/review. In **E**, no symbols were reported given that the best final model did not include the factor condition. Instead, FDR-corrected p-values were provided. In **A-E**, dots represent individual subjects’ performance.

A more detailed analysis revealed that subjects not only reviewed more often for difficult trials than for easy ones, but also that decisions were related to categorization accuracy (Figure 2C). Reviews were significantly more frequent after an incorrect response on the first choice than after a correct one (mixed-effect *glm*, interaction feedback x difficulty, F=13.3, p=1.04e-5). The percent of reviews after a correct choice decreased with easier conditions, but concomitantly increased after incorrect choices. The difference incorrect versus correct decreased with increased difficulty. Thus, subjects were able to detect their own errors and reviewed appropriately, and this ability was greater when confronted with simple categorizations. This indicates that uncertainty about one’s own performance was higher in the most difficult condition, inducing more decisions to review.

Response times can reflect the process by which confidence contribute to the decision to review. Indeed, subjects were significantly slower to choose when they subsequently reviewed than when they subsequently validated (Figure 2D, mixed-effect *glm*, factor decision, F=684.9, p=7.4e-143). Subjects were slower when making an incorrect choice compared to a correct one, independently of the subsequent decision (validate or review) and more strongly for difficult trials (interaction feedback x difficulty, F=9.37, p=8.6e-5; Wald test, p<0.03 for all incorrect/correct comparisons. No other interactions survived model selection). At decision time (review or confirm), subjects were also slower following incorrect choices compared to correct ones, and slower when reviewing compared to confirming their choice (Figure 2E, mixed-effect *glm*, factor decision, F=63.4, p=2e-15; factor feedback, F=8.51, p=3.5e-3). Difficulty as well as interactions did not survive in that case. Even though similar effects were observed for both response times, differences appeared substantially greater at the choice stage (right/left target selection) than the decision stage. Such observations revealed that subjects’ confidence continuously impacted behaviour at all stages and even before the appearance of decision targets.

### Lack of behavioural adaptation following a review in Humans

The above results mostly show that subjects use the review option in an adapted manner. However, the main goal of metacognitive abilities is not only to estimate how uncertain subjects are in a given situation, but also to promote information seeking to improve performance. To investigate such process, we calculated the categorization sensitivity (type1) and metacognitive sensitivity (type2) for each subject and for the three different conditions (see Methods for details). As expected, type1 sensitivities decreased with difficulty (mixed-effect *glm*, F=87.9, p=5.5e-15) (Figure 3A). However, type2 sensitivities revealed an unexpected result. Even though type2 sensitivities varied significantly with difficulty (mixed-effect *glm*, F=3.96, p=0.028), subjects showed relatively low metacognitive abilities, with type2 sensitivities close to 0 in the most difficult conditions. Also, type1 and type2 sensitivities were significantly correlated (Spearman linear correlation, R=0.401, p=0.011, Figure 3B). This suggests that subjects’ metacognitive performance depended partially on sources of variation that affected their categorization performances. This might have been the case if for instance subjects used the review option to discriminate difficult vs. easy trials instead of interring the correctness of their choice independently of the categorization’s difficulty.

**Figure 3.**
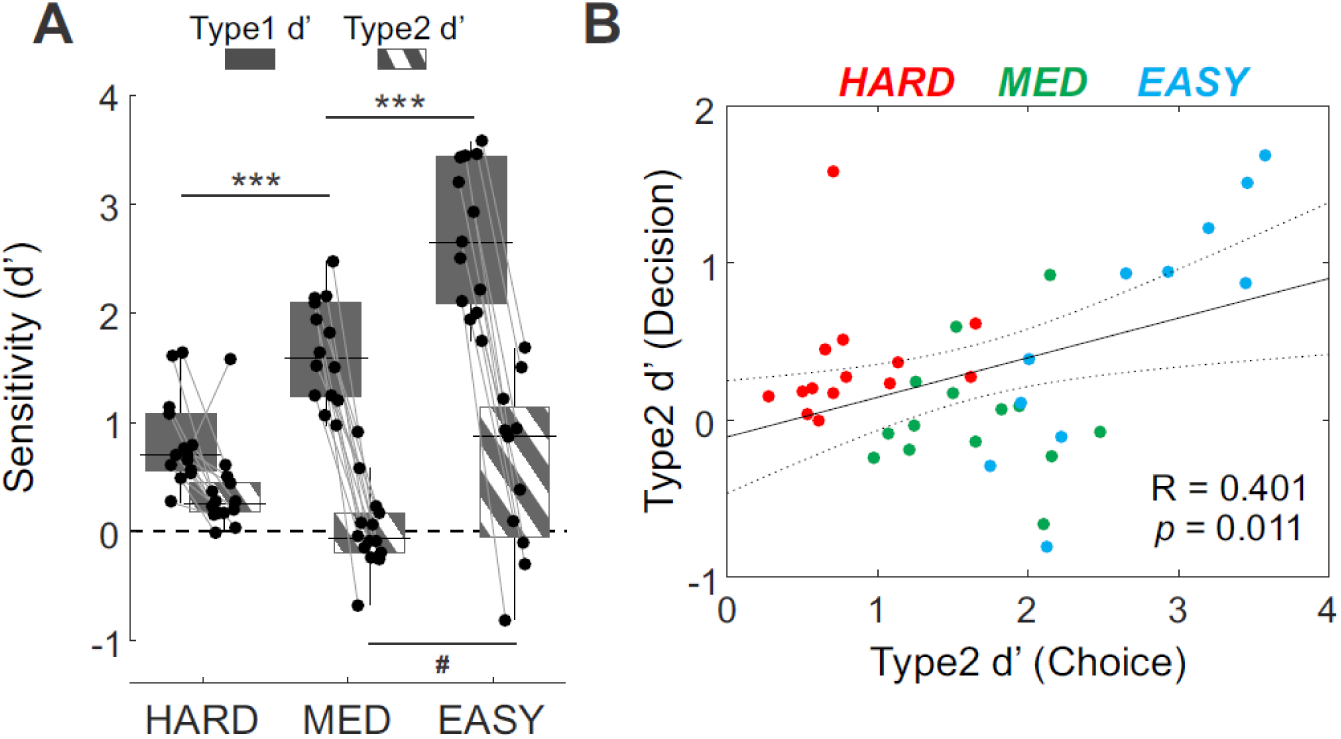
Choice and Decision sensitivities. **A**, Average sensitivity measures (d’) for Type1 (Choice, plain grey) and Type2 (Decision, grey stripes) ROC depending on difficulties. Significant post-hoc comparisons (Wald test with FDR correction) are reported as follow: ***p<0.001 for type1 comparisons; #p<0.05 for type2 comparisons. **B**, Individual d’ for type2 plotted against type1, for each difficulty level (Hard in red, Medium in green, Easy in blue). The dotted line represents the Pearson’s linear correlation (±95% CI), independent of difficulty.

Even if subjects showed markers of their ability to detect errors, they were relatively unable to correct them (as suggested in Figure 2B). Consistent with the overall low level of metacognitive sensitivity, further analysis revealed unexpected consequences of the review process. First, subjects showed on average a significant bias toward repeating the same right/left choice after a review (i.e. percent of shift across difficulties was below 50%, Wilcoxon sign rank test, z=-2.54, p=0.011) (Figure 4A), and this bias toward confirmation was not modulated between conditions (mixed-effect *glm*, factor difficulty: F=1.97, p=0.152). Figure 4B presents performance of subjects at different levels of review and depending on performance: in grey the performance for first validated choices (D1, no review); in red, final performance following an incorrect first choice (whatever the number of reviews, one or more), and in blue, the final performance following a correct first choice. Note that in the HARD condition, the final performance was particularly low when subjects initially made a mistake (i.e. red bar for HARD, Wald test p<0.001) (mixed-effect *glm*, interaction difficulty x Trial type, F=2.74, p=0.03). Similar decreases in performance following incorrect trials compared to correct ones was also observed for MED and EASY conditions, albeit to a lesser extent (post-hoc comparison, p<0.018). Alternatively, no differences were observed between the immediately validated performance (D1) compared to the final performance following a correct choice in the first selection (COR) (Wald test, all conditions at p>0.39). Thus, the review process worsened performance only when the trial started with a mistake. In other word, confusion and/or confirmation biases arose after errors during the successive reviews.

**Figure 4.**
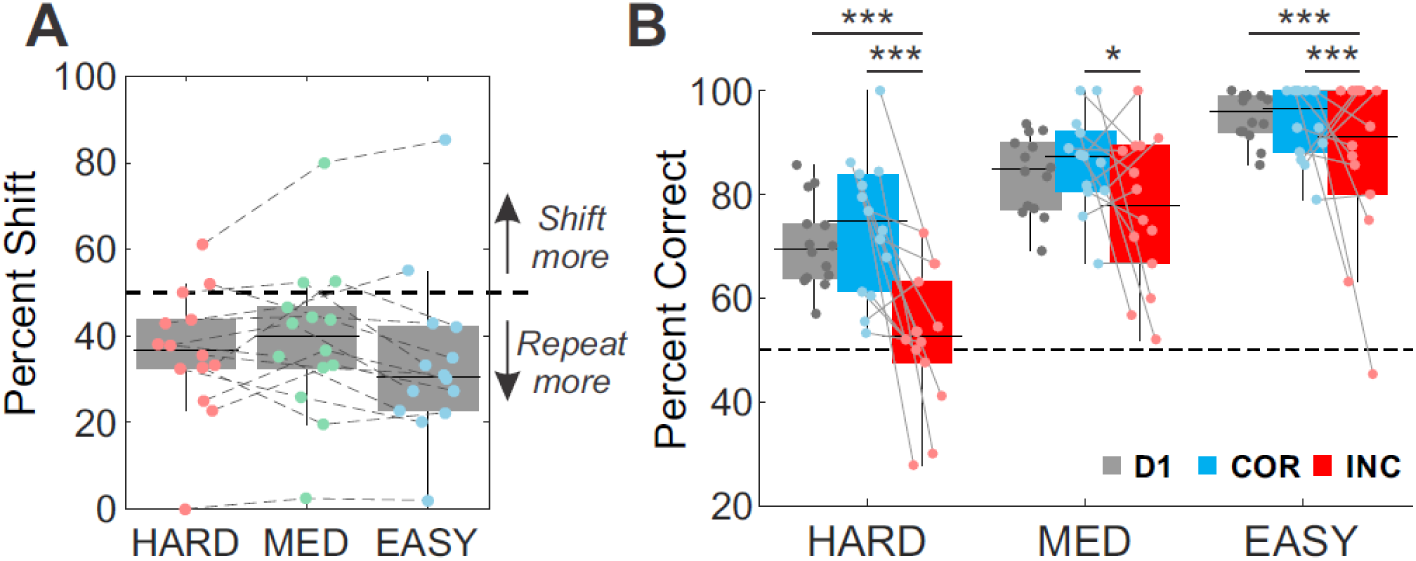
Performance changes following a review. **A**, Average percent of shift in response compared to the previous choice following a review. **B**, Average performance before (grey, first Choice response) and after reviews (blue and red, final Choice response). Final performance was separated depending on the first choice (correct first choice in blue and incorrect first choice in red). Significant post-hoc comparisons (Wald test with FDR correction) are reported as follow: *p<0.05, **p<0.01, ***p<0.001.

### Categorization performance in monkeys

Two male rhesus monkeys were tested in a task similar to the one used for human subjects (Figure 1A, **bottom panel**). Monkeys behaved correctly in the categorization test, by showing appropriate psychometric performance. The discrimination thresholds differed between monkeys, with monkey H being more accurate with lower angles than monkey D (Figure 1C). To elicit 60, 70 and 85% of correct responses, monkey H was tested with bar orientations of 1°, 2° and 5° relative to the vertical (5°, 10° and 20° for monkey D). Both monkeys had slightly better performance for rightward stimuli compared to leftward (0.15° and 3.8° of performance difference between right and left orientations for monkey H and D respectively).

To assess whether reaction and movement times varied between conditions, we used a mixed-effect *glm* for each individual monkey and measure (choice RT, choice MT, decision RT, decision MT), with a random-effect of sessions on the intercept. All initial models included the 3 following factors: feedback, difficulty and decision, as well as all possible interactions. The results for the best models are reported below (see also Methods for model selection procedure). Choice RT were slower when monkeys made an incorrect choice compared to a correct one (mixed-effect *glm*, factor feedback, monkey D: F=40.1, p=2.9e-10; monkey H: F=14.6, p=1.3e-4; factor difficulty and decision, as well as interactions did not survive model selection). Decision RT also increased following incorrect choices for both monkeys, with an additional interaction with difficulty (mixed-effect *glm*, interaction feedback x difficulty, monkey D: F=4.5, p=0.011; monkey H: F=7.3, p=6.8e-4) (Wald test, all difficulty p<0.04, except for monkey H in the hard condition, p=0.33). In only one monkey (monkey D) choice MTs were slower following incorrect compared to correct trials, especially in Hard and Easy conditions (mixed-effect *glm*, interaction feedback x difficulty, F=3.13, p=0.04) (Wald test, p<0.024, but not in Med condition, p=0.08). Decision MT in monkey D did not depend on feedback or difficulty, but significantly change with the subsequent decision to review or validate, as reported in the next section. Overall, monkeys tended to be slower to plan their choice when incorrect, but this was mostly independent of the difficulty of the categorization.

### A sub-optimal use of the review option in monkeys

After an initial use of the review option during the first sessions, both monkeys stopped doing so for a long period of time (Figure 5A). Such behaviour forced us to slightly change the task to familiarize them with the review option. Two main changes were tested: forced review trials were included (i.e. in which the validate option was not available) and correct/incorrect feedback was also adjusted to increase review benefits (i.e. more juice reward when correct and less time penalty when incorrect after a review, these parameters also changed over time) (see also “Notes on training procedure with monkeys” in *Methods*). Contrary to our predictions, these changes did not elicit voluntary reviews (Figure 5A, arrows ‘a’). A final change allowed monkeys only one single review, without requiring the validation of their last choice. This was done at session number 61 and 112 for monkey D and H respectively (arrows ‘b’ in Figure 5A). This modification was efficient in the sense that both monkeys freely selected the review option again after a few sessions, without resorting to a heavier training approach. Thereafter, their behaviour remained stable. For the purpose of this study, we focused our analysis on sessions where monkeys used the review option in a stable way, from session number 64 and 120 until the end for monkey D and H respectively (*n* = 22 & 38 sessions for monkey D and H) (Figure 5A, highlighted in grey).

**Figure 5.**
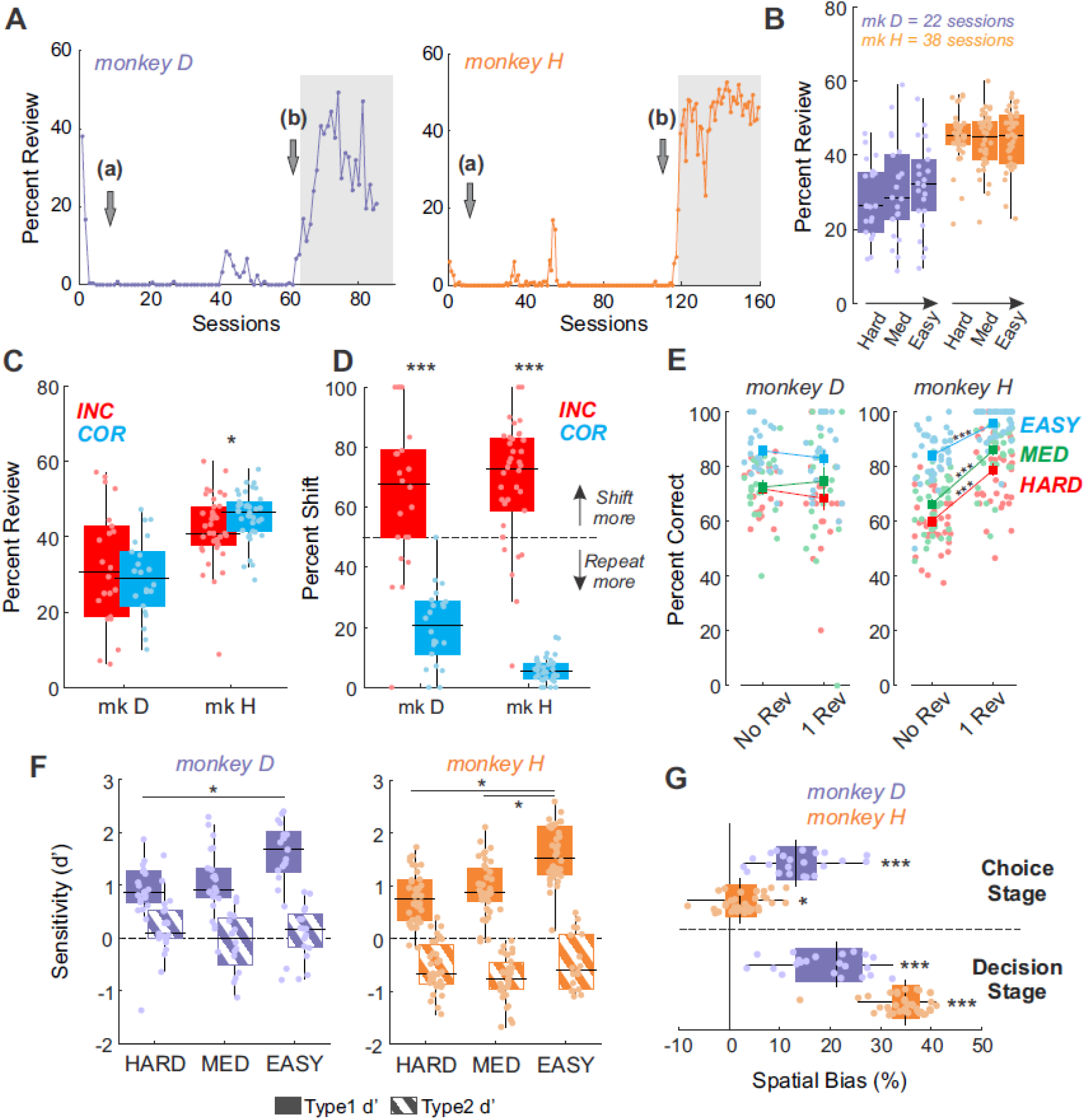
Behavioural performance of monkeys. **A.** Percent of reviews over sessions for monkey D (left panel) and H (right panel). Grey arrows indicate major changes in task design: (a) forced reviews were introduced; (b) the first session where monkeys were proposed to review only once, without the need to validate the second choice. Grey highlight represents the sessions included in the following analyses. **B & C**, Percent of reviews depending on conditions (**B**) and previous performance (**C**; correct in blue; incorrect in red). **D**, Percent of shifts depending on previous performance. **E**, Final performance when monkeys validated immediately (No review) or after 1 review depending on conditions (Hard in red; Medium in green; Easy in blue) and for both monkeys. **F**, Average sensitivity measures (d’) for Type1 (Choice, plain) and Type2 (Decision, stripes) ROC depending on difficulties. **G**, Spatial bias (tendency to choose a particular side rather than a particular target) for Choice (Right or Left) and Decision (Review or Validate) stages for both monkeys. Significant post-hoc comparisons (Wald test with FDR correction) are reported as follow: *p<0.05, **p<0.01, ***p<0.001. Dots in **B-G** represent individual sessions’ performance.

Contrary to what was observed with human subjects, monkeys did not use the review option differently with different levels of difficulty (Figure 5B) (mixed-effect *glm*, factor difficulty; monkey D: F=2.05, p=0.13; monkey H: F=0.41, p=0.66). Moreover, if monkeys were using the review option based on their own uncertainty, one would expect a greater proportion of reviews following incorrect choices compared to correct ones, as observed in human subjects. This was not the case (Figure 5C). In fact, an opposite effect was observed for monkey H (mixed-effect *glm*, factor feedback; monkey D: F=1.25, p=0.27; monkey H: F=5.86, p=0.017). However, both monkeys repeated more their choices following correct choices compared to incorrect ones, a situation where they usually shifted (mixed-effect *glm*, factor feedback; monkey D: F=57.1, p=2.3e-9; monkey H: F=372, p=1.3e-30, Figure 5D). Such observation suggests a potential benefit of the review process.

As described in the previous section, we did not observe clear modulations in RT and MT depending on the subsequent decision to review or validate. Only decision MT in monkey D were slower for confirmed choice compared to reviewed ones (mixed-effect *glm*, factor decision, F=19.8, p=8.7e-6). In all other cases, the factor ‘decision’ did not survive model selections. The absence of RT/MT modulations suggests both monkeys might not have use the review option as an expression of their uncertainty.

The behavioural benefits of the review process in terms of performance were different in the two monkeys (Figure 5E). Specifically, for monkey D reviewing was neither advantageous nor deleterious, the performance being modulated only by the difficulty (mixed-effect *glm*, monkey D, factor difficulty: F=12.3, p=1.3e-5; no other factor or interaction survived model selection). However, monkey H showed a significant increase in performance on reviewed trials, modulated by the difficulty (mixed-effect *glm*, monkey H, interaction difficulty x nbReview, F=4.77, p=9.3e-3; post-hoc comparison for each difficulty, Wald test p<4.5e-17). The discrepancy between monkeys might be explained by differences in experimental conditions. Monkey D was allowed to see the stimulus longer during the review (+400ms, representing a duration gain of 80% on average), whereas monkey H was proposed a slightly simpler stimulus after session number 94 (+3°, representing an ideal performance gain of 16.25% on average). Such differences tends to suggest that reviewing the same stimulus (as in monkey D) did not increase performance at all, even when displayed longer.

Similar to what we found in human subjects, monkeys’ type1 sensitivities decreased with difficulty (Figure 5F) (mixed-effect *glm*, factor difficulty, monkey D: F=4.05, p=0.022; monkey H: F=18.2, p=1.4e-7; post-hoc comparisons, Wald test, monkey D: Hard vs Easy, p=0.027; other comparisons, p>0.051; monkey H: all comparisons at p<1.4e-5, except Hard vs Med at p=0.31). This was not the case for type2 sensitivities, where low values were observed as well as no modulation (mixed-effect *glm*, factor difficulty, monkey D: F=1.98, p=0.15; monkey H: F=2.58, p=0.08).

In order to avoid spatial biases, but also to limit preparatory responses, we randomized both Right/Left targets and Review/Validate levers locations between trials (on the right or left of the screen). Yet we observed that both monkeys developed a strong spatial bias toward one side, especially at the decision stage (Figure 5G) (Wilcoxon sign-rank test, choice stage; monkey D: z=4.1, p=4e-5; monkey H: z=2.24, p=0.025; decision stage; monkey D: z=4.1, p=4e-5; monkey H: z=5.37, p=7.7e-8). The use of this low cognitively demanding strategy (spatial) strongly suggests that monkeys poorly discriminated between review or validate options.

Taken together, our findings reveal that monkeys were unable to develop and use the opportunity to review or validate the test in an optimal manner. Despite a long and careful training, no clear evidence supported a metacognitive evaluation at the decision stage in monkeys. It is important to acknowledge the possibility that a heavier training approach might have been able to elicit such behaviour. This was however not the objective as we were trying to obtain a natural development of a review strategy.

## Discussion

In this study, we designed a protocol to assess confidence judgments under uncertainty, but also the resulting behavioural adaptations. Inspired by information-seeking tasks, this novel metacognitive task used reviews as a means to reveal confidence and was intended to test both human and non-human primates. Observations in human subjects performing this task was in accordance with previously reported results, showing that subject might be able to report confidence with the opportunity to review or validate. However, in monkeys, the behavioural study did not reveal expected review vs. validate behaviour. The main issue might be that, in both species, reviewing did not improve performance and rather competed with, or was perturbed, by alternative low-cognitive cost strategies. In this context, the design might have hinder monkeys to use the review option appropriately.

The behavioural task induced a review process triggered by difficulty and estimated performance. Subjects’ uncertainty on choice was reflected in the use of this option. They showed behavioural markers of error detection, even before any feedback was delivered. This confirmed previous observations that human subjects can adequately report retrospectively their confidence in a perceptual choice^3^, although our protocol reveals subjects’ confidence through the measure of review behaviour and does not require explicit report. In this sense, this protocol might be useful from a clinical perspective, especially when trying to understand why obsessive compulsive disorder (OCD) patients showed excessive checking behaviour and impairments in self-performance monitoring^18,19^. Moreover, from a neurophysiological perspective, studies of performance monitoring generally investigate adaptive processes after subjects receive feedback on performance^20–22^ (but see ^23^). Our protocol allows us to study markers of performance monitoring in the absence of external feedback.

Metacognitive processes might also serve to promote correction of putative mistakes, in order to improve decisions^1,17^. Apparently, the current task design did not allow subjects to improve their performance after the review process. Rather, subjects developed a bias toward repeating the same choice even if they mostly reviewed after incorrect choices. Such confirmation bias, often reported in the literature^24^, might contribute to the absence of improvement. But it might also reveal that subjects used reviews as a form of verification, with the objective of reducing their uncertainty about their first decision. This is in contrast to the use of review to simply revise their first choice (as it would be the case for changes of mind). In that sense, our task design might well capture the use of metacognitive evaluation for adaptive change in control in order to acquire more information. Nevertheless, given that both underlying reasons might contribute to subjects’ use of reviews, further experiments are needed to reveal subject’s actual strategies at the trial level, for example by varying and controlling the *quantity* of information given at any moment. Using appropriate modelling tools and experimental changes, it might be possible to dissociate between subjects’ overall bias (i.e. reviewing only for confirmation about an uncertain first choice) and decisions that depend on recently collected information.

The absence of improvement following a review might also explain why the signal detection approach indicated poor metacognitive abilities. In the procedure, type2 measures reflect the ability to detect and correct wrong decisions^25^, and scores of this measure were very low. This should not be taken as the sole measure of metacognition. Subjects could not adequately adapt their behaviour under high uncertainty (i.e. following a review), but had chosen to review in a way that indicates some metacognitive process. This dissociation therefore underlines the difficulty of finding a single measure of metacognition, and the fact that several factors impinge on this ability. One interpretation is that, in the present context, the review process counterintuitively generates interference or doubts that prevent the addition of information across the successive reviews and hence the increase in performance. After the first choice, during review, the perceptive information that is offered again to the subjects appeared not to be integrated and cumulated. Rather, subjects might have based their decisions on purely internal information related to performance (the perception of an initial error) or memory of the first perceived stimulus. In this context reviews are detrimental. A study with OCD patients similarly observed that patients seemed to use mnemonic cues to respond to a discrimination test after numerous reviews and did not use the available perceptual information anymore^26^. More recently, other experiments report similar detrimental effect of reviews in healthy subjects^27–29^. We argue here that subjects did so especially when uncertainty was maximal (in the more difficult condition), by making a choice depending only on a poor memory of the stimulus and a pure guess on the current correct response. If this appears to be true, we might expect changes in neurophysiological markers that are incongruent with the information provided by the stimulus.

Finally, the existence of a correlation between perceptual and metacognitive abilities suggests that both rely on the same underlying estimation (i.e. perceptual in this case). The difference in choice response times between reviewed trials and validated ones also support such interpretation. Nevertheless, relationships between perceptual and metacognitive measures have been often reported in the literature (for example ^4,30,31^ but see ^2^), and have been considered as an argument toward a metacognitive explanation of human and non-human strategies^5,7^. In our case, subjects might decide whether to review or validate a choice depending on the speed of their responses, by assuming that a long deliberation might be related to greater uncertainty^6^. Even if not metacognitive in the sense defined by Nelson & Narens^1^, this gives rise to a widely accepted proposition that metacognitive abilities are based on an *indirect* access to one’s own cognition, by making inferences from observable cognitive processing results^6,7^. Specifically, ease of processing, retrieval fluency or cue familiarity might be used to report confidence and have been argued to reflect metacognitive processes^4,5,32^. The present data are in favour of such theories.

Taken together, our observations highlight that the task might be appropriate to study metacognition from an information-seeking point of view. Review behaviour allowed us to address two metacognitive mechanisms: the monitoring of decisions and the related adaptation that might occur. However, subjects’ behaviour suggests that the review option was mostly for self-confirmation rather than a way to modify a choice. Post-decision adaptation was sub-optimal in our subjects, possibly due to task design issues as discussed next.

Recent studies have shown the possibility to test some forms of checking behaviours in monkeys^33,34^. In the present study, we were not able to elicit reviews in monkeys that would be based on their perceived uncertainty. Despite weak evidence of their ability to adequately use the review option, monkeys quickly fell into a non-optimal and cognitively less-demanding strategy to perform the task. Such failure might be explained by different factors as discussed below.

First, the difficulty to elicit reviews in monkeys might depend on the learning procedure we adopted, or a related issue concerning the task design itself. Assessing metacognitive abilities in monkeys requires avoidance of the development of alternative strategies^35^. In particular, reward-induced biases or external cue associations need to be tightly controlled so as to avoid confounds with the intended effect of metacognitive process. In the literature, many studies were debated due to the possible use of alternative non-metacognitive processes^8,9,13,35^. We tested whether a protocol with an adaptive review process would be efficient. However, as observed by Son & Kornell^11^, even simpler protocols by comparison to the one we developed, using high and low-bets following a perceptual response, are tricky to use in monkeys. In their study, authors reported their failure to elicit an efficient use of the decision stage, due to the expression of a bias toward a specific option (selecting only the high-bet option). However, after a long training and many modifications in the task design, they were able to elicit what they consider to be an optimal use of bet options^11,12^.

In our task, the credit assignment problem, i.e. figuring out the link between a specific choice (right or left) and a delayed reward (after the decision stage), was arguably the most challenging element. Credit assignment is a complex issue especially in sequentially structured tasks or in multiple choice situations^36^. For monkeys, one way to bypass such an issue might be to always validate a choice and not take into account the decision stage. This was our monkeys’ first strategy. Importantly, when we introduced the single review option, monkeys soon used the opportunity to review. In this case, the association between choice and feedback was sometimes present (after a review, the feedback was given immediately after the second right/left choice), and monkeys changed their strategies accordingly. However, both monkeys also adopted a simpler spatial strategy. Reviewing might have been perceived as effortful, and as similar as cancelling a previous choice. Even if a visual cue indicated trial transitions, adding contextual information to clarify the structure of a trial might have helped (e.g. changing the background colour from one trial to another).

Another possible explanation was that monkeys never perceived the benefits associated with the review option, even if analyses revealed an advantage for one individual. Reviewing a choice underlies a greater cost than validating, in terms of physical and cognitive effort at least. Delay and effort are two separate features that both depreciate human and non-human decisions when experienced^37,38^. Even if the review vs. validate options were equalized in duration, our results might suggest that our monkeys were more sensitive to effort than delay, and so were less willing to review a choice than to wait during the time penalty after an error.

Finally, one might question the ability of monkeys to use their own confidence. The failure to induce reviews in our experiment is certainly not only explained by a natural inability to express metacognition. Nevertheless, this question is hotly debated in the literature^32,39,40^. Information-seeking protocols appear to be highly relevant to study metacognitive processes in animals as one-trial tests have revealed abilities of self-knowledge based adaptive behaviour in macaques^16^. Importantly, macaques showed appropriate patterns of responses in uncertainty test whereas new world monkeys (capuchin) did not^41^. Yet, few studies reported the relative volatility of metacognitive responses or even the absence of such response in many individuals and different species tested with or without extensive training^11,14,42^. Concomitantly, two main criticisms have been proposed against the existence of metacognition in non-human animals. The first is that simple heuristics like reward associations, and not metacognition, guide behaviour in information-seeking or uncertainty response paradigms^10,17,35,40^. The second questions the relevance of metacognition to address problems to which animals are confronted^39^. Even if animal research tried to answer these issues^43^, more investigations and new protocols are required to confirm whether and how non-human animals have access to metacognition^32,44^.

Few improvements are required to adapt our protocol to the study of metacognitive evaluation and control in animals for behavioural and neurophysiological studies. One possibility to explain the failure to induce reviews in monkeys is that, like in humans, the transfer of information from one trial to the next during the review process was not optimal. Hence the review was not beneficial. If this is the case, adding contextual information to link decisions and outcomes across actions and events might help. Similarly, information transfer might not have been facilitated by using fixed stimuli like oriented gratings. Dynamic stimuli (random dot motion for example) could contribute to enhance the gain of perceptual information after a review, and so might be beneficial.

## Methods

### Participants and Apparatus

#### Humans

Fourteen subjects (7 males and 7 females, aged 20-34 years, mean = 24.15 years) participated in this study after giving informed consent. The study was carried out in accordance with the recommendations of the Code de la Santé Publique and performed in a laboratory approved by the “Agence Nationale de Sécurité des Médicaments et des produits de santé (ANSM)”. All subjects had normal or corrected-to-normal vision and all were right-handed. Testing was performed on an Apple computer (Apple Inc., USA) using Matlab (MathWorks Inc., USA) and the Psychtoolbox^45^. Subjects were comfortably seated 19.6 inches (50cm) away from a 23-inch screen, on which visual stimuli were displayed. Responses were made by using arrows on a computer keyboard with their right (dominant) hand. Experiments were done in a low-luminosity room.

#### Monkeys

Two male rhesus monkeys (*Macaca mulatta*), weighing 8 kg and 12 kg (monkeys D & H respectively) were used in this study. All procedures followed the European Community Council Directive (2010) (Ministère de l’Agriculture et de la Forêt, Commission nationale de l’expérimentation animale) and were approved by the local ethical committee (CELYNE, C2EA #42, project reference: C2EA42-13-02--0402-10). Monkeys were trained to perform the task while seated in a primate chair (Crist Instrument Co., USA) in front of a tangent touch-screen monitor (Microtouch System, Methuen, USA). An open-window in front of the chair allowed them to use their preferred hand to interact with the screen (monkey D, left-handed; monkey H, right-handed). The position and accuracy of each touch was recorded on a computer, which also controlled the presentation of visual stimuli via the monitor (CORTEX software, NIMH Laboratory of Neuropsychology, Bethesda, MD). During experiments, monkeys were not head-restrained and eye movements were not controlled.

### Behavioural Task

#### Humans

Subjects were first asked to perform a categorization task, based on the orientation of a stimulus following a staircase procedure to adjust difficulty (Figure 1A, **top panel**). Each trial started with a 1000ms fixation period during which subjects fixated a central dot. Then, a stimulus, consisting of a low-contrast Gabor patch oriented from vertical reference either on the left or on the right, was presented centrally on a grey background during 200ms. After a 500ms delay, subjects reported the orientation of the stimulus by using the right or left arrow keys of a standard keyboard. After reporting their choice (right or left) and an additional delay of 1000ms, they were asked to either validate or review (re-execute) the test (with either the bottom or the top arrow keys respectively). If subjects decided to validate their choice, a visual feedback was displayed centrally for 800ms, consisting of the word “correct” (shown in green) or “incorrect” (in red), and a new stimulus was presented on the subsequent trial. A review was potentially triggered by a subjective lack of confidence. If subjects decided to review, the same stimulus was presented again and a new choice could be made. Note that following half of the reviews, subjects were presented with a longer stimulus than on its first presentation, for a duration of 250ms instead of 200ms. The trial ended only after the subject validated their choice. Correct trials were not rewarded *per se*, but incorrect trials were penalized by a time penalty of 15 seconds. A 1000ms delay was introduced between trials.

Subjects were required to perform 420 trials, divided into 6 blocks of 70 trials (note that 1 subject performed blocks of 65 trials instead). Between blocks, subjects were able to take rest. At the start of the experiment, instructions were given to explain the nature of the task as described above. Emphasis was placed on the general idea that the review option could help the subject complete the experiment more quickly, i.e. that reviews and consequent improvement of performance would compensate for the time-out after errors.

#### Monkeys

Monkeys were trained to perform a task similar to the one use with humans, where they were required to perform a categorization test based on the orientation of a bar, and then to either validate or review their choice (Figure 1A, **bottom panel**).

To initiate a trial, monkeys had to touch and hold a lever item, represented by a grey triangle on the bottom of the screen. Once touched, a central dot appeared on the screen for 800ms. Then, the central dot turned off and the stimulus appeared on the screen (a grey rightward or leftward oriented bar) for a duration between 400 and 900ms (this duration was fixed during a session but changed across training). After a delay of 200ms, two oriented bars (one oriented 45° to the left, the other 45° to the right relative to the vertical) were used as targets. The relative position of targets was randomized from one trial to another (e.g. the leftward bar might be either positioned on the right part or the left part of the screen, randomly). Monkeys reported their choice by touching one of the targets. This was followed by an additional delay of 200ms. The two decision levers were then displayed, allowing monkeys to review or validate their previous choice. The review option was represented by a grey inverted triangle lever, and the validate option by a grey disk. The position of each lever on the screen (bottom right or left) was randomly assigned from one trial to the next.

If monkeys touched the validate option they received a feedback corresponding to their performance: correct choices were rewarded by a squirt of apple juice lasting between 300 and 1000ms, incorrect choices were penalized by a grey screen lasting between 10000 and 15000ms (note that rewards and penalties changed over the course of training, hence the range of values). To equalize review/validate options duration, the duration of a penalty for incorrect validation was set to be equal to two review trials. However, it is important to keep in mind that, by design, review trials are more effortful than validated ones, given the number of touches required. After the feedback delivery, a visual signal was displayed on the screen, consisting of a red circle lasting 800ms, and indicating the change of condition.

If monkeys touched the review option, the central dot appeared again on the screen for 800ms and the stimulus was displayed. Two particular modifications of the task were used to stimulate the review process. First, the duration of the stimulus was increased by 400ms after each review. In some of monkey H’s sessions, the duration of the stimulus was not increased, but instead the stimulus became easier (larger angle) after reviews (this was the case for all the sessions analysed thereafter, see Results for details). The following events were the same as described above. Second, feedback duration was modified if monkeys reviewed at least once. Correct choices were more rewarded after a review than after no review, with duration of reward between 500 and 1400ms (+91.6% and +77.7% of reward for monkey D and H respectively). Also, incorrect choices were less penalized after reviews, with a time penalty between 1000 and 3000ms (−82.6% and −79.4% for monkey D and H respectively). These explicit benefits of the review were introduced to help monkeys during the training procedure (by increasing the review utility), but were not intended to be used after the completion of the training (see below for details on training).

### Staircase & Psychometric analysis

To maintain different levels of uncertainty during the categorization, three different stimulus orientations were used randomly across trials (i.e. HARD, MED and EASY).

For human subjects, orientations were determined depending on subject’s own performance using a classical staircase procedure prior to the experiment. In this procedure (which lasted 240 trials), subjects had to choose whether the stimulus was leftward or rightward oriented, without the possibility to review or validate their choice. Stimulus orientations were defined depending on subjects’ performance, with 3 randomly mixed staircase rules^46^ (one-up one-down, one-up two-down and one-up three-down). The use of the 3 parallel staircases procedure (80 trials each) allowed us to assess subjects’ performance more accurately. Based on performance during the staircase procedure, we calculated a psychometric curve for each subject, using a binomial generalized linear model (logistic regression). Three absolute stimuli orientations were then extracted and used during the main experiment to induce 70%, 75% and 85% of correct responses (HARD, MED and EASY conditions respectively, see Figure 1B for an average psychometric curve and related orientations).

Similarly, for monkeys, orientations were not fixed between sessions but varied depending on their performance in preceding sessions (by considering the 5 to 10 last sessions). We computed a psychometric curve from past monkeys’ performance using a binomial generalized linear model (logistic regression) (Figure 1C). Then, stimuli orientations were selected to elicit 70, 80 or 90% of correct responses on average. Such procedure allowed us to maintain uncertainty in categorization trials across sessions independently of other learning-related processes.

### Notes on training procedures with monkeys

We first trained monkeys on the categorization task alone, which lasted several months. During this period, performance feedback was given immediately after their choices. Once performance in the categorization task was stable and sufficiently accurate, monkeys were then introduced to the decision stage, allowing to review or validate choices. This was done without any particular shaping. That is, we did not imposed any familiarisation with one of the two decision stage options prior to this stage, initially to avoid as much as possible biases toward a specific behaviour (review or validate). Over-training monkeys could lead to non-optimal strategies and is particularly problematic when it comes to understand natural behaviour such as monkeys’ metacognitive ability^16^. However, during the learning phase with the decision stage, we faced a few issues that led us to modify the task design in several steps. Specifically, after experiencing the review option both monkeys stopped using it within a few sessions (see Figure 5A). We thus implemented three main changes: 1) a small proportion of forced review trials (10-25%) were introduced (on these trials, only the review target is presented at the first decision stage), 2) penalties and rewards were adjusted throughout the training to increase the utility of the review and 3) monkeys could not review more than one time in a row, without the need to validate after the review (i.e. no second decision stage were proposed, and importantly, feedback was delivered immediately after the second choice).

Here, we report behavioural data for 22 and 38 sessions (for monkey D and H respectively). Sessions were selected based on the frequency in choosing the review option. To accurately assess putative metacognitive performances, we excluded sessions before the monkeys selected the review option in a stable manner (see Results for details).

### Behavioural Analysis

#### Humans

Classical performance measures were calculated for each subject. Specifically, choice response times (i.e. time between the appearance of the right/left targets and the button press) and decision response times (i.e. time between the appearance of the review/validate targets and the button press) were recorded and analysed. Response times exceeding 10 seconds were rejected from further analyses (n=3 and 1 trials for choice and decision times, respectively, across all subjects). To improve the normality of the data distribution, we used the log transform of response times as dependent variable in the statistical analyses. Also, we calculated the percentage of review decisions (whatever the number of successive reviews), depending on conditions or depending on previous performance (correct or incorrect in the first selection). Initial and final performance was computed for each trial. Moreover, subjects’ strategy following a review was studied by calculating the percent of shift, which is the proportion of reviewed trials after which the initial choice (before review) was changed in the decision that was made after reviews. As very few subjects made more than four successive reviews, these trials were entirely removed from further analyses (n=6/5850 trials). For similar reasons, we restricted our analyses of the performance against the number of review (Figure 2B) as well as type1/type2 performance (Figure 3) to a maximum of two successive reviews (n=5731/5844 included trials, 2% of trials were rejected).

As mentioned previously, the stimulus was displayed for an extra 50ms following half of subjects’ reviews. This manipulation did not elicit any difference in performance, as assessed using mixed effect models (see Statistical procedure for more details). Specifically, the performance after 1 review was only modulated by difficulty; the factor Stim Length did not survive model selection. For the performance after 2 successive reviews, we found a significant interaction (mixed effect *glm*, interaction difficulty x Stim Length, F=3.51, p=0.035) but post-hoc comparisons revealed no statistically significant differences between stimulus length levels (Wald test, W<1.48 and p>0.42 for all comparisons). For this reason, all trials were pooled together in further analyses, independently of stimulus length.

#### Monkeys

In addition to the measures used for human subjects, choice reaction times (choice RT; i.e. time between the appearance of oriented targets and the release of the lever) and choice movement times (choice MT; i.e. time of arm movements from lever release to the touch on selected target) were computed on each trial. Decision reaction times (decision RT; i.e. time between the appearance of the decision levers and the release of the target) and movement times (decision MT; time between the target release and the touch of a decision lever) were also recorded. As for human subject analyses, we used the log transform of the reaction and movement times as dependent variables in statistical analyses. Moreover, a measure of spatial bias was calculated for both stages (choice and decision) by dividing the number of touches made ipsilateral to the hand used by the total number of touches (note that monkey D and H use their left and right hand respectively).

### Signal Detection Theory approach

To assess subjects’ performance in the categorization test and their metacognitive abilities, we used the signal detection theory approach implemented by MacMillan & Creelman^25^. For this purpose, we computed the receiver operating characteristic (ROC) curve for both categorization (type1) and metacognitive (type2) performance. In this analysis, all trials are designated a ‘trial type’ amongst HIT, Correct Rejection (CR), False Alarm (FA) and MISS. In a 2-choice task such as ours, a reference for this classification must be chosen. It is the reference that differentiates type1 and type2 performance.

For type1 we sought to measure categorization performance, and the reference was trials on which the stimulus was oriented to the right. Hence a trial that had a rightward stimulus and a response ‘right’ was classified as a HIT, whereas a trial that had a leftward stimulus and a response ‘left’ was classified as CR. It follows that rightward stimulus and response ‘left’ was Miss, and leftward stimulus and response ‘right’ was a false alarm. This referencing approach allowed us to complete a ROC analysis despite the two-choice nature of the task.

In the type2 analysis, we sought to measure the metacognitive performance, i.e. whether the subject correctly self-diagnosed their performance in their choice to review or not. Hence the reference was trials where the response was both correct AND validated. In this case a HIT was a correct and validated choice, whereas a CR was an incorrect then reviewed response. As such, FA was a correct reviewed response, whereas a MISS was an incorrect validated response.

Finally, we took into consideration the number of reviews (none, one or two) made within a trial, because this can be considered as a rating scale of certainty - no review meant that subjects were sure, 2 reviews that they were really unsure of their response. Hence, for a given trial, each new choice after a review or 2 reviews was assigned a cumulative probability of HIT and FA, based not only on the outcome of that choice but also of the previous choices within the same trial. We then derived ROC curves from cumulative HIT and FA probabilities before applying a Gaussian z transformation (i.e. zROC). We fitted a linear curve in order to obtain the slope of the zROC. We further defined d’_1_ and d’_2_, the horizontal and vertical intercept of the zROC curve, respectively:

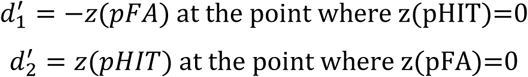

From there, we then extracted the sensitivity measures (type1 and type2 d’_a_) according to the formula **[1]** defined in Chapter 3 of MacMillan & Creelman’s guide^25^:

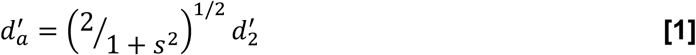

where *d’*_*a*_ represents the sensitivity and *s* the slope of the linear zROC curve. This measure of sensitivity, contrary to the classic d’, is able to characterize a ROC curve using a single value of distance from each pair of HIT and FA probabilities.

### Statistical procedures

All statistical procedures were performed using Matlab. Behavioural differences between conditions were assessed using mixed-effect generalized linear models (*glm*). For human subjects, each model included a random-effect on the intercept of the variable that the model was fitting (e.g. performance, percent of review, or the log of response times), to account for baseline differences between subjects (random-effect of subjects). For monkeys, models included a random-effect on the intercept to account for baseline differences between sessions (random-effect of sessions). Across models, the following factors (and levels) were considered: Difficulty (Hard, Medium, Easy), Feedback (Incorrect, Correct), Number of Review (0 to 2), Decision (Review, Validate), Trial Type (Validated choice, Correct choice Reviewed, Incorrect choice Reviewed), and Stimulus Length (post-review increased or not). For every model used, we first performed a model selection by repeatedly testing the effect of dropping the least significant factor (starting with the interactions). The most appropriate models were then selected using Log Likelihood ratio test (with p<0.05). F-tests for each fixed-effects term in the selected models were reported in the Results section. Post-hoc comparisons were performed by computing the estimated marginal means^47^ and using Wald test. P-values were corrected with False Discovery Rate (FDR) to account for multiple comparisons.

## Acknowledgements

We thank K. N’Diaye and L. Mallet for helping with the task design and for their comments on the results. We thank J. Benistant for the help in the acquisition of data from human subjects. We also thank M. Valdebenito, M. Seon and B. Beneyton for animal care and C. Nay for administrative support. This work was supported by the Agence National de la Recherche (DECCA ANR-10-SVSE4-1441), the Fondation pour la Recherche Medicale (FMS and DEQ20160334905) and was performed within the framework of the LabEx CORTEX (ANR-11-LABX-0042) of Université de Lyon, within the program “Investissements d’Avenir” (ANR-11-IDEX-0007). E.P is employed by the Centre National de la Recherche Scientifique.

## Data availability

All relevant data and codes are available from the authors upon request.

## Notes

**Conflict of Interest:** No. Authors report no conflict of interest

